# Generation and characterization of a *Dkk4-Cre* knock-in mouse line

**DOI:** 10.1101/2023.05.23.541881

**Authors:** Houda Khatif, Hisham Bazzi

## Abstract

Ectodermal appendages in mammals, such as teeth, mammary glands, sweat glands and hair follicles, are generated during embryogenesis through a series of mesenchymal-epithelial interactions. Canonical Wnt signaling and its inhibitors are implicated in the early steps of ectodermal appendage development and patterning. To study the activation dynamics of the Wnt target and inhibitor *Dickkopf4* (*Dkk4*) in ectodermal appendages, we used CRSIPR/Cas9 to generate a *Dkk4-Cre* knock-in mouse (*Mus musculus*) line, where the *Cre* recombinase cDNA replaces the expression of endogenous *Dkk4*. Using *Cre* reporters, the *Dkk4-Cre* activity was evident at the prospective sites of ectodermal appendages, overlapping with the *Dkk4* mRNA expression. Unexpectedly, a predominantly mesenchymal cell population in the embryo posterior also showed *Dkk4-Cre* activity. Lineage-tracing suggested that these cells are likely derived from a few *Dkk4-Cre*-expressing cells in the epiblast at early gastrulation. Finally, our analyses of *Dkk4-Cre*-expressing cells in developing hair follicle epithelial placodes revealed intra- and inter-placodal cellular heterogeneity, supporting emerging data on the positional and transcriptional variability in placodes. Collectively, we propose the new *Dkk4-Cre* knock-in mouse line as a suitable model to study Wnt and DKK4 inhibitor dynamics in early mouse development and ectodermal appendage morphogenesis.

## Introduction

Ectodermal appendage morphogenesis is initiated through a series of reciprocal interactions between the mesenchyme and the overlying epithelium (Hardy, 1992). The molecular cross-talk leads to the formation of histologically visible local thickenings of the epithelium called placodes that give rise to teeth, nails, mammary glands, sweat glands and hair follicles (HFs) (Pispa & Thesleff, 2003). How these placodes form and are uniformly patterned, are still largely open questions in animal biology.

HFs are arranged in a regularly-spaced pattern in the skin that has been proposed to arise through a reaction-diffusion mechanism, with an interplay between inducing and inhibitory signals originating from a common source (Turing, 1990). Essential to this model are multiple signaling pathways like the canonical Wnt pathway, which is an early inducer in the HF developmental hierarchy (Andl et al., 2002; Glover et al., 2017).

Wnt signaling is activated upon the binding of Wnt-ligands to their cognate receptors Frizzled and Low-density Lipoprotein Receptor*-*related Protein (LRP) 5/6 co-receptors. Ultimately, the Wnt signal is transduced through the stabilization and nuclear translocation of β-catenin, which together with the lymphoid enhancer factor/T-cell factor (Lef/Tcf), initiate the transcription of Wnt target genes (Nusse & Clevers, 2017). To prevent the constitutive accumulation of β-catenin, Wnt is antagonized by a set of inhibitors, including the secreted proteins from the Dickkopf (DKK) family 1-4 that bind to the LRP5/6 co-receptors, leading to the assembly of the β-catenin destruction complex (Mao et al., 2001; Zorn, 2001). Together with the transmembrane protein Kremen (KREM), DKKs and LRP6 form a ternary complex, which ensures the removal of LRP6 from the cell surface, thereby inhibiting Wnt activity (Mao et al., 2002).

In this context, DKK4 has been shown to synergistically act with KREM to exert its Wnt inhibitory functions (Patel et al., 2018). *Dkk4* is also a canonical Wnt target and proposed to act in a negative feed-back loop with Wnt signaling during ectodermal appendage morphogenesis (Bazzi et al., 2007; Sick et al., 2006). Moreover, stratified epithelia-specific overexpression of *Dkk4,* whose mRNA is normally restricted to ectodermal placodes (Bazzi et al., 2007; Sick et al., 2006), affects secondary HF development (Cui et al., 2010; Sima et al., 2016), and decreases the number of sweat glands (Cui et al., 2014). Studying this interplay between Wnt and DKK4 during development is important to understand how negative feedback loops pattern tissues and organs.

At the initiation of gastrulation, the posterior part of the epiblast undergoes an epithelial-to-mesenchymal transition at the primitive streak, whose location is determined by Wnt activity (Mohamed et al., 2004). The gastrulation movements organize the germ layers, where the epiblast cells that do not ingress through the primitive streak adopt an ectodermal fate, and the pre-somitic mesoderm develops in the posterior region (Bardot & Hadjantonakis, 2020). A hallmark of organogenesis is the formation of the somites that derive from the pre-somitic mesoderm and develop sequentially to give rise to the future bones, muscles, connective tissue and mesenchyme including a large part of the skin dermis (Tam, 1981). However, compared to Wnt signaling, which has been extensively studied during development and in other contexts, the regulation of *Dkk4* expression in early mouse (*Mus musculus*) development warrants further investigation and requires the generation of new models and tools.

In this study, we describe the generation of a *Dkk4-Cre* knock-in mouse line to investigate *Dkk4* transcriptional regulation in response to Wnt signaling activity during early development and in ectodermal appendage formation, particularly in HFs. We use lineage-tracing of the *Dkk4-Cre-*reporters to show that the reporter is active in ectodermal appendages. Our data also reveal that a handful of cells are induced in the epiblast around the initiation of gastrulation and likely give rise to posterior somites and mesenchymal cells in the posterior region of the embryo. Notably, our data also suggest cellular heterogeneity within and among the HF placodes.

## Results

### Ectodermal appendages are marked by *Dkk4-Cre*-expressing cells

To study the Wnt-DKK4 signaling interplay in ectodermal appendage formation, we used CRISPR/Cas9 to engineer a *Cre*-recombinase knock-in mouse line that utilizes the *Dkk4* endogenous promoter. The *Cre* cDNA was successfully inserted into the start codon of exon1 in the *Dkk4*-locus and to disrupt and replace the expression of the *Dkk4* open reading frame (ORF) (**Figure 1a, b and Supplementary Figure 1**). To assess the *Cre*-recombinase expression, we crossed heterozygote *Dkk4-Cre* mice with a dual-reporter line that expresses *tdTomato* or *EGFP* at the cell membrane (*mTmG*) (Muzumdar et al., 2007), or in the nucleus (*nTnG*) (Prigge et al., 2013). Upon induction of the *Cre* recombinase in *Dkk4*-expressing cells, the *tdTomato* gene, which is flanked by two *loxP* sites, is excised and *EGFP* is expressed instead (**Figure 1c**). Thus, the cells that are derived from *Dkk4*-*Cre*-expressing progenitors will maintain the *EGFP* signal and their lineage can be traced over time. To test the specificity of the *Dkk4-Cre* line, we investigated the expression of *EGFP* at ectodermal appendages, where *Dkk4* mRNA has been shown to be expressed using *in situ* hybridization (**Figure 2**) (Bazzi et al., 2007; Sick et al., 2006). EGFP-positive cells localized at the prospective sites of the primordia of the dental field (embryonic day (E)10.5), vibrissae follicles (E12.5), mammary glands (E13.5), pelage or back-skin HFs (E14.5) and eccrine glands on the palmo-plantar skin (E18.5), but not cranial placodes (data not shown), confirming that the activity of the *Cre* recapitulated the *Dkk4* mRNA expression (**Figure 2a, b and Table 1**). At E18.5, we also observed EGFP-positive signals in the heart (**Supplementary Figure 2a**), kidney (white arrow, **Supplementary Figure 2b**), adrenal gland (yellow arrow, **Supplementary Figure 2b**) and other organs including muscles (data not shown), indicating that *Dkk4*-expressing cells contribute to other tissues and organs besides ectodermal appendages.

**Figure 1:**
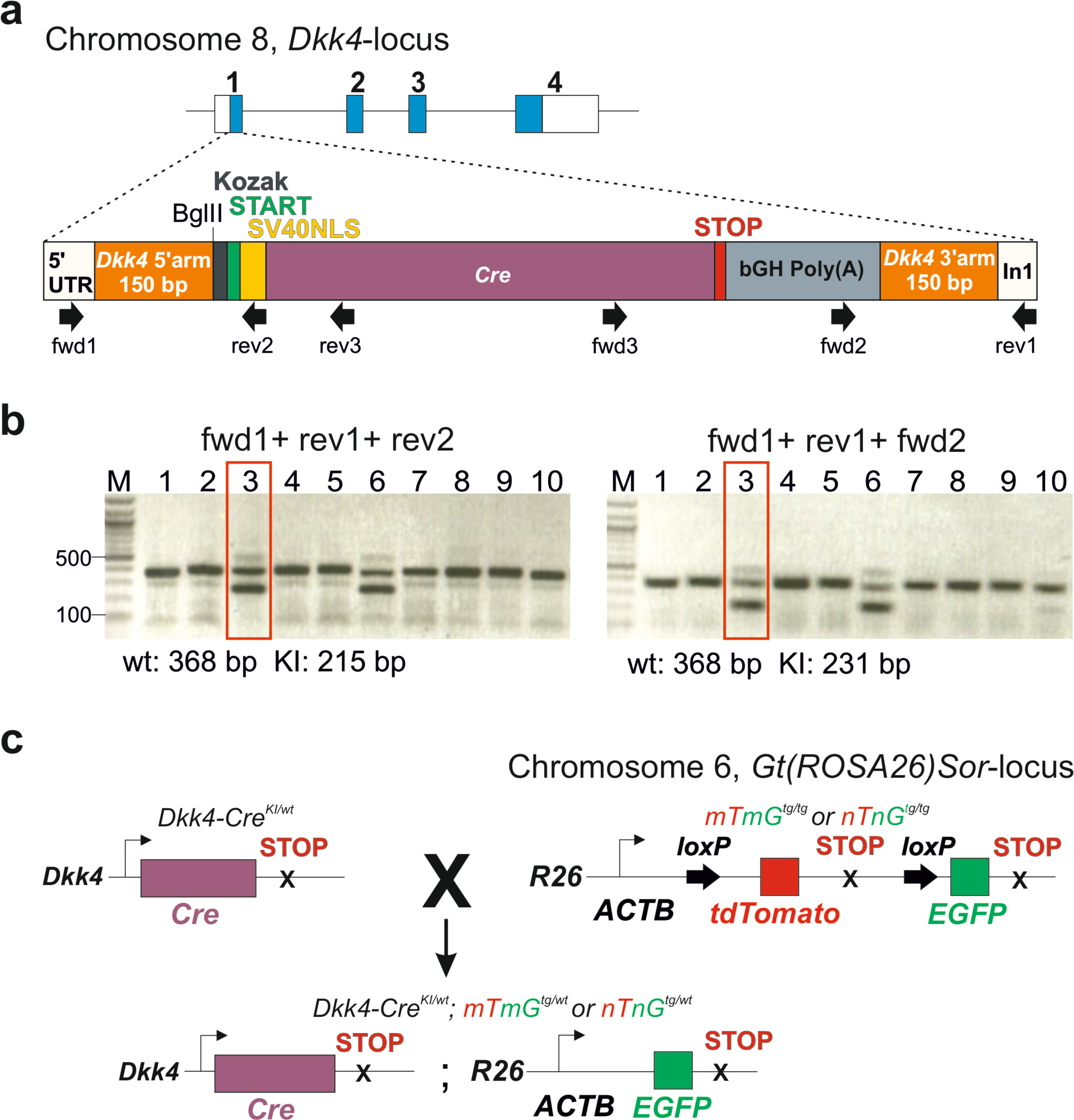
*Cre* insertion into the *Dkk4*-locus using CRISPR/Cas9. (a) A schematic representation of single-stranded (ss) DNA template of ∼1.6 Kb encoding *Cre* cDNA, a nuclear localization signal (NLS) of the Simian Virus (SV) 40 T antigen and a bovine growth hormone polyadenylation signal (bGH Poly(A)). *Cre* was inserted into Exon1 of *Dkk4* via homology-directed repair with arms of homology of 150 bp. The primers used for genotyping and sequencing are depicted below (fwd and rev). 5’ untranslated region, UTR and Intron1, In1. (b) PCR analyses of the wild-type (wt) and knock-in (KI) alleles. The 368 bp PCR-product from (fwd1+rev1) corresponds to the wt band (top band in Founder 3, for example). The *Cre* insertion was verified at the 5’ end by the 215 bp KI-band (fwd1+rev2, bottom band in Founder 3 on the left), and at the 3’ end by the 231 bp KI-band (fwd2+rev1, bottom band in Founder 3 on the right), as represented in (a), and the products were confirmed by sequencing (**Supplementary Figure 1**). Founder 3, red box, was used for further breeding and experiments. (c) Scheme of *Dkk4-Cre^KI/wt^* male mice crossed to *mTmG^tg/tg^* or *nTnG^tg/tg^* females. The *mTmG/nTnG* alleles are at the ROSA26 (R26) locus and driven by the chicken beta-actin core promoter (ACTB) (Muzumdar et al., 2007; Prigge et al., 2013). The tdTomato gene is flanked by *LoxP* sites and excised in cells that express *Cre*. The embryonic progeny with the genotype *Dkk4-Cre^KI/wt^;mTmG^tg/wt^* or *Dkk4-Cre^KI/wt^;nTnG^tg/wt^*and EGFP-positive expression were used for lineage-tracing analyses.

**Figure 2:**
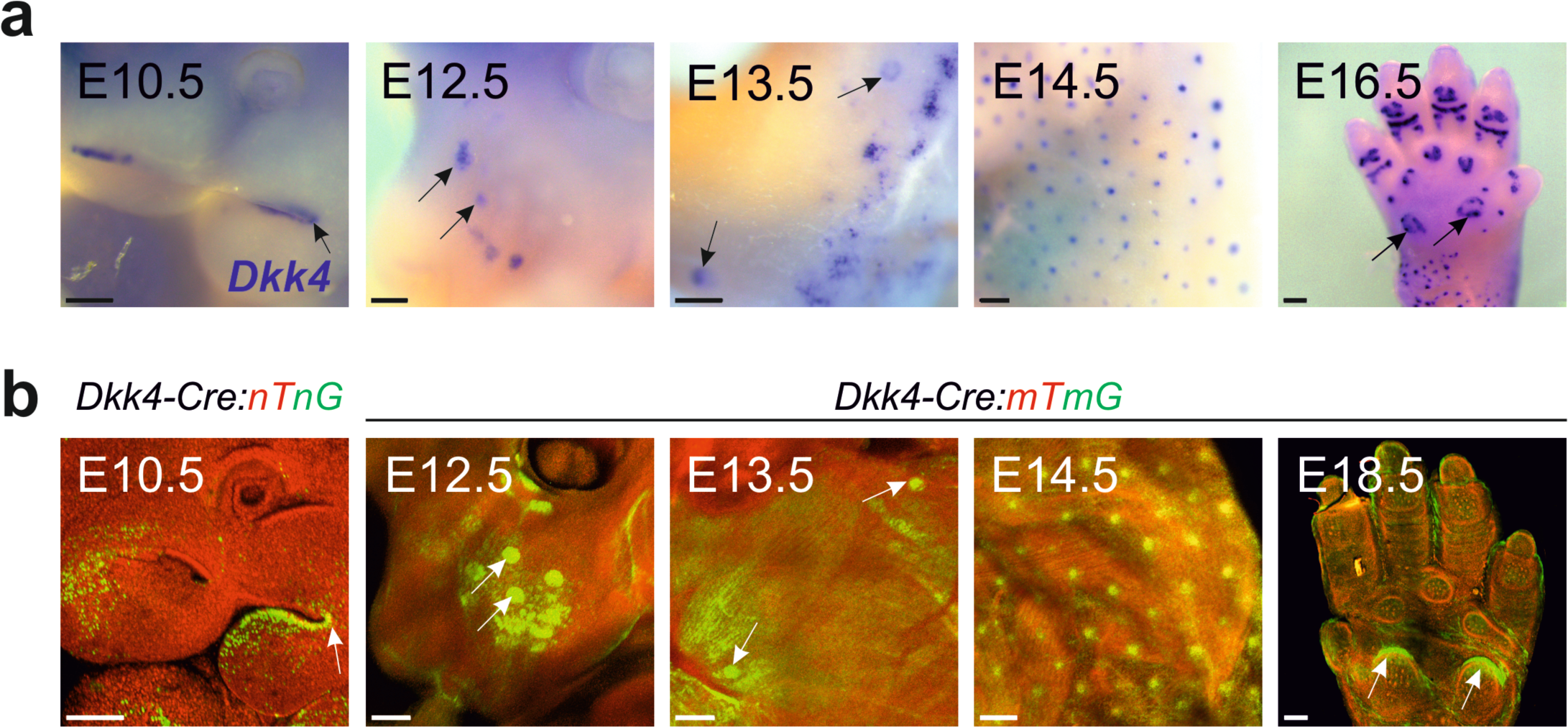
*Dkk4* mRNA and *Dkk4-Cre-*induced EGFP-reporter expression in ectodermal appendages. (a) Whole-mount *in situ* hybridization of *Dkk4* mRNA localization on wt embryos at different developmental stages in the presumptive dental field (arrow at E10.5), vibrissae follicle placodes (arrows at E12.5), mammary gland placodes (arrows at E13.5), back-skin hair follicle placodes (E14.5) and eccrine glands (arrows at E16.5) (scale bars: 200 µm). (b) Confocal images of whole-mounts of *Dkk4-Cre^KI/wt^:nTnG^tg/wt^*(E10.5) or *Dkk4-Cre^KI/wt^;mTmG^tg/wt^* (E12.5, E13.5, E14.5, E18.5) embryos showed a similar EGFP-expression pattern to the *Dkk4* mRNA in (a) (scale bar: 200 µm). Anterior is up in E10.5 – E14.5.

**Table 1:** Summary of the *Dkk4* mRNA compared to *Dkk4-Cre*-reporter expression in mouse embryos. The *Dkk4* mRNA expression was assessed using colorimetric-based whole-mount *in situ* hybridization, and the *Dkk4-Cre* was analysed using *nTnG* or *mTmG* reporters.

| Region | *Dkk4* mRNA | *Dkk4-Cre* |
| --- | --- | --- |
| Epiblast/ Primitive streak | unknown | yes |
| Pre-somitic mesoderm | no | yes |
| Somites | no | yes |
| Dental field | yes | yes |
| Vibrissae follicles placodes | yes | yes |
| Mammary gland placodes | yes | yes |
| Hair follicle placodes | yes | yes |
| Eccrine glands | yes | yes |
| Dermis | no | yes |
| Muscle | no | yes |
| Heart | unknown | yes |
| Adrenal glands | unknown | yes |

### A subset of *Dkk4*-*Cre*-expressing cells arises around gastrulation

In addition to the expression of the *Dkk4-Cre* in ectodermal appendages, lineage-tracing in *Dkk4-Cre* reporter embryos suggested that a developmentally earlier cell population expressed *Dkk4-Cre* (**Figure 3**). Specifically, 90% of dissected *Dkk4-Cre^KI/wt^*-reporter embryos showed evident EGFP-positive cells in the posterior region between E6.5 and E15.5 (**Figure 3a, b**), while the remaining 10% exhibited a rather wide-spread distribution of EGFP-positive cells (**Supplementary Figure 3a**). Because of the mostly consistent pattern of the EGFP signals and given that *Dkk4 in situ* hybridization did not show *Dkk4* mRNA expression in the same region in early embryos (E8.5 and beyond, data not shown), we reasoned that the progenitors of these posterior cells arose earlier during development. In agreement, lineage-tracing revealed a few EGFP-positive cells in the epiblast of early gastrulating mouse embryos at E6.5 and E7.5, some of which co-localized with the primitive streak and mesoderm marker Brachyury (**Figure 3b and Supplementary Figure 3b**), and correlated with increased EGFP-positive cells in the pre-somitic mesoderm at E8.5 (**Figure 3b, right**), as well as in the posterior somites between E9.5-E10.5 (**Figure 3a, right**). Sagittal sections of *Dkk4-Cre:nTnG* embryos at E14.5, with predominantly posterior expression, indicated that in the anterior region, EGFP-positive cells were mainly found in the epithelial skin layer and mostly localized to the HF placodes (**Figure 3c**). However, in the posterior region, the EGFP-positive cells were also found in the dermis (**Figure 3c**), likely representing the same lineage of the earlier mesenchymal population. In addition, a section through the skin and subcutis in the posterior region at E18.5, revealed EGFP-positive cells in the connective tissue layers, muscle as well as in the dermis and epidermis (**Figure 3d and Table 1**).

**Figure 3:**
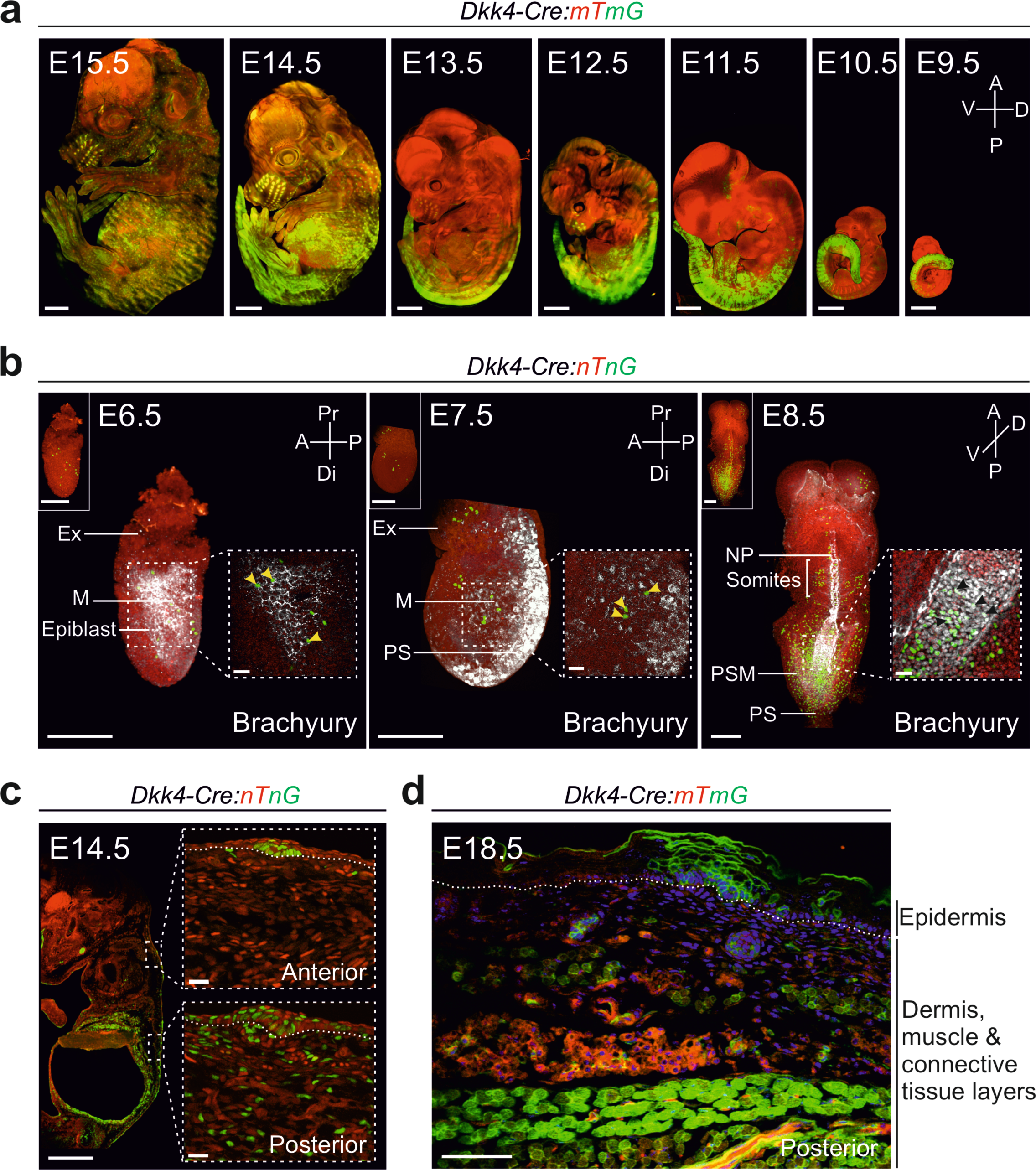
*Dkk4*-*Cre-*expressing cells are induced in the early epiblast and spread to the mesenchymal cells of the posterior embryo region. (a) Confocal images of E9.5 – E15.5 (right to left) *Dkk4-Cre^KI/wt^:mTmG^tg/wt^* embryos with evident EGFP-expression in the posterior body region and hair follicles (scale bar: 1000 µm). Anterior (A) is up, posterior (P) is down, ventral (V) is to the left and dorsal (D) is to the right. (b) Confocal images of E6.5 – E8.5 *Dkk4-Cre^KI/wt^:nTnG^tg/wt^* embryos that were immune-stained with Brachyury (scale bar: 200 µm). Dashed squares: single confocal Z-planes show co-localization of some EGFP-positive cells (yellow arrowheads) and Brachyury in the “wings of mesoderm (M)” (scale bar: 30 µm). In E6.5 and E7.5 embryos, anterior (A) is to the left, while the posterior (P) region, where the primitive streak (PS) is localized, is to the right. The proximal (Pr) region is up and marked by the extraembryonic region (Ex), whereas the distal (Di) region is down. E8.5 embryos with a ventral (V) to dorsal (D) view are oriented along the anterior (A) – posterior (P) axis, with the notochordal plate (NP), the pre-somitic mesoderm (PSM) and somites shown. (c) Sagittal section of an E14.5 *Dkk4-Cre^KI/wt^;nTnG^tg/wt^* embryo (scale bar: 1000 µm) with anterior or posterior dorsal skin regions enlarged and rotated 90° counter-clockwise in the insets (scale bar: 20 µm). (d) Sagittal section of an E18.5 *Dkk4-Cre^KI/wt^;mTmG^tg/wt^* embryo showing the different skin layers expressing EGFP at the posterior embryo region (scale bar: 50 µm). A dotted line demarcating the basement membrane between the epidermis and dermis is shown in (c) and (d).

### The hair follicle epithelium is not solely derived from *Dkk4-Cre*-expressing cells

Given the wide-spread expression of the *Dkk4-Cre* lineage in the posterior region of most embryos, we focused on the anterior skin region, which showed more restricted epithelial expression that appeared to arise later in development. In particular, we assessed *Dkk4-Cre* reporter expression in developing HFs, as a representative of ectodermal appendages. Interestingly, EGFP-positive cells were detected in the basal layer of E12.5 and E13.5 epidermis, even before any morphological signs of HF placode formation (**Figure 4a**). As expected at E14.5 and E15.5, the EGFP-positive cells were found in HF placodes (yellow arrows in **Figure 4a**), but also occasionally in patches of the inter-follicular epidermis (white arrows in **Figure 4a**). Intriguingly, a closer inspection of the *Dkk4-Cre* reporter expression in HF placodes revealed that not all placodal cells were EGFP-positive, suggesting that a subset of cells in the placode did not descend from *Dkk4-Cre*-expressing cells. Co-staining with the placode markers LHX2 or SOX9 showed that, on average, around half of the LHX2- or SOX9-expressing placodal cells were EGFP-negative (arrows in **Figure 4b, c**). In fact, the wide distribution of the double positive cells indicated intra- and inter-placodal heterogeneity (**Figure 4c**). In confirmation of these findings, in growing HFs at post-natal day (P) 8, only a fraction of cells in the matrix and HF layers seemed to be derived from *Dkk4- Cre* reporter-expressing cells (**Figure 4d**). The data suggest a heterogeneous origin of the HF epithelium that includes and goes beyond the *Dkk4-Cre* reporter-expressing cells in the placode.

**Figure 4:**
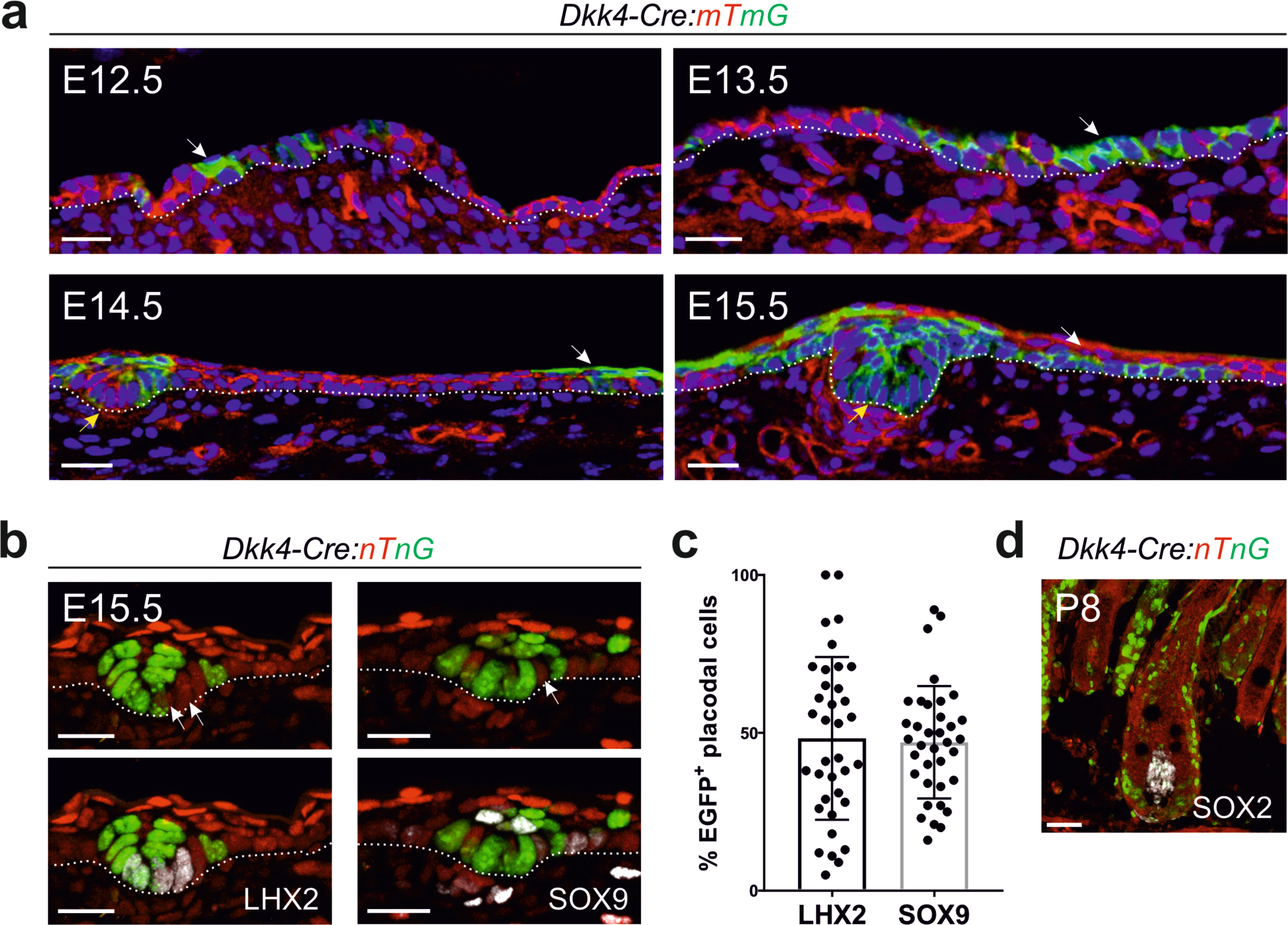
*Dkk4-Cre-*derived EGFP-positive cells mark a subset of hair follicle placodal cells and inter-follicular epidermis. (a) Sections from the anterior back-skin of E12.5 – E15.5 *Dkk4- Cre^KI/wt^;mTmG^tg/wt^* embryos, showing EGFP-expression in the epidermis (white arrows at E12.5- E13.5), the inter-follicular epidermis (white arrows at E14.5 – E15.5) and hair follicle placodes (yellow arrows at E14.5 and E15.5) (scale bar: 20 µm). (b) Sagittal sections of hair placodes from the anterior region of E15.5 *Dkk4-Cre^KI/wt^;nTnG^tg/wt^* were stained with the placode markers LHX2 or SOX9 that co-localized with ∼50% of EGFP-positive cells (scale bar: 20 µm). A dotted line demarcating the basement membrane between the epidermis and dermis is shown in (a) and (b). (c) Quantification of the percentage of EGFP*-* and LHX2*-* or EGFP*-* and SOX9-expressing placodal cells shown in (b) from the anterior back-skin region, where every dot represents one hair placode from a total of 6 embryos each. (d) Sagittal section of a post-natal hair follicle from a P8 *Dkk4-Cre^KI/wt^;nTnG^tg/wt^* back-skin immuno-stained with SOX2 to mark the dermal papilla (scale bar: 50 µm).

## Discussion

We describe the generation of a *Dkk4-Cre* knock-in mouse line that reproduces the expression of *Dkk4* mRNA in ectodermal appendages (Bazzi et al., 2007), using *Cre*-reporter lines. Unexpectedly, the lineage-tracing strategy uncovered a rare population of individual cells around early gastrulation (E6.5 – E7.5) that expresses *Dkk4-Cre*. The progeny of these cells mostly occupies the pre-somitic mesoderm (E8.5) and subsequently the posterior somites (E9.5), likely giving rise to the mesoderm-derived tissues in the posterior region of developing embryos from E10.5 – E18.5. In addition, the data suggest that these cells contribute mostly to mesodermally-derived organs such as the heart and kidney. Some of the single *Dkk4-Cre*-expressing cells at E6.5 – E7.5 co-localized with the primitive streak and wings of mesoderm marker Brachyury (Wilkinson et al., 1990). Interestingly, Wnt is first activated in pre-gastrulating E6.5 embryos (Maretto et al., 2003; Mohamed et al., 2004), covering the *Dkk4-Cre* expression domain. *Dkk4* is a direct target gene of canonical Wnt signaling and is hypothesized to act in a negative feedback loop to fine tune Wnt activity in ectodermal appendages (Bazzi et al., 2007; Sick et al., 2006). The site of the primitive streak and the induction of the mesoderm is largely dictated and coordinated by the canonical Wnt pathway (Mohamed et al., 2004). Therefore, it is highly likely that the early *Dkk4*-*Cre*-expressing cells at E6.5 are induced by high Wnt activity in these cells. One caveat of our knock-in/ knockout strategy is that reducing the dose of DKK4 in heterozygous *Dkk4-Cre* embryos is expected to reduce the inhibition on canonical Wnt signaling, likely potentiating the Wnt pathway and enhancing the expression of *Dkk4-Cre*, which itself is a target of Wnt, resulting in ectopic *Cre* expression in this rare cell population and perhaps others in later development (see below).

The ectoderm and its lineages are derived from the anterior epiblast, which do not express *Dkk4- Cre* around the beginnings of gastrulation (E6.5 – E7.5). This finding suggest that the vast majority of ectodermal *Dkk4-Cre*-driven reporter expression arises later in embryogenesis coinciding with the *Dkk4* mRNA and is predominantly in the placodes of developing ectodermal appendages, including HFs (Bazzi et al., 2007; Sick et al., 2006). The *Dkk4-Cre*-driven reporter expression in the inter-follicular epidermis between E12.5 and E14.5 suggests that the *Dkk4* expression in the skin epithelium is not restricted to the placode as initially suggested by colorimetric *in situ* hybridization, a technique with insufficient resolution to definitively ascertain single-cell expression.

We also observed that not all cells in the HF placode expressed *Dkk4-Cre*. The basal cells of the placode are labelled by LHX2 expression (Rhee et al., 2006), while the SOX9-positive cells mostly occupy the suprabasal part and periphery of the placode (Morita et al., 2021; Nowak et al., 2008; Ouspenskaia et al., 2016). In vibrissae HFs, Morita et al. suggested the existence of four genetically distinct concentric layers that form the HF placode and give rise to the different compartments of the mature HF (Morita et al., 2021). The *Dkk4* mRNA was mainly detected in the center of vibrissae HF placodes, while *Sox9* localized at the periphery, similar to our observations for the back-skin (Morita et al., 2021). The heterogeneity of the *Dkk4*-*Cre*-driven reporter expression in the HF placode is also reflected in later stages of HF morphogenesis, with cells in the matrix and HF layers labelled in a mosaic manner. Our data suggest heterogeneity in the placodal cell fates that give rise to the different parts of the HF epithelium, as has been recently shown by time-lapse imaging and single-cell-RNA-sequencing (Morita et al., 2021). Collectively, the *Dkk4-Cre* mouse line provides a new tool to study Wnt signaling induction of the DKK4 inhibitor in early gastrulation and ectodermal appendage morphogenesis, using lineage-tracing and conditional knockouts.

## Methods

### Generation of Dkk4-Cre^KI/wt^ mice

The CRISPR/Cas9-mediated *Dkk4-Cre* knock-in mice (*Dkk4^tm1(Cre)Baz^*) were generated by the CECAD *in vivo* Research Facility (ivRF, Branko Zevnik) using pronuclear microinjection of the gRNA (*5’-AGAGUGACUGAGGAUGGUAC-3’*), Cas9 mRNA, Cas9 protein and a single stranded DNA (ssDNA) template into fertilized zygotes of FVB/NRj mice (Troder et al., 2018). The approximately 1.6 Kb ssDNA shown in **Figure 1a** was ordered from Integrated DNA Technologies, Inc. (IDT; Coralville, IA, USA). Through homology directed repair and homologous recombination, the *Cre* cDNA was inserted into and replaced the *Dkk4* ORF (location of insertion: Chr8:23,114,199 before the ATG of Exon 1, www.ensembl.org). The single-stranded template DNA harbored *Dkk4* homology arms of 150 bp, a BglII restriction site, a Kozak sequence, SV40NLS and bGH Poly(A). The mice were genotyped using PCR for the inserted *Cre* DNA at the 5’-region (*forward1*, *fwd1: 5’-CGGAAGAGTATGCTGGTCAG-3’* and *reverse2*, *rev2:* 5’- *GCCATGCCCAAGAAGAAGAG-3’)* and 3’-region *(fwd2: 5’-TGGGAAGACAATAGCAGGCA-3’ and rev1: 5’- GCGCCCTCACCTCTAAGT-3’*). The primers rev3 (*5’- CGAGTGATGAGGTTCGCAAG-3’*) and fwd3 (*5’-CGCTGGAGTTTCAATACCGG-3’*) were used for sequencing. Only *Dkk4-Cre* male mice were used for experimental breedings. Phenotypes were analyzed in the FVB/NRj; C57/BL6NRj mixed background (*Dkk4*-*Cre^KIIwt^*; *mTmG^tg/wt^*or *nTnG^tg/wt^*). The mTmG (Jax Labs stock #007576) (Muzumdar et al., 2007) and nTnG (Jax Labs stock #023035) (Prigge et al., 2013) mice are available from the Jackson Laboratory (Jax Labs; Bar Harbor, ME, USA), but were back-crossed to the FVB/NRj and C57/BL6NRj backgrounds in this study. The animals were housed and bred under standard conditions in the CECAD ivRF under a 12 h light cycle, at a temperature of 22 ± 2°C, 55 ± 5% relative humidity and with food and water *ad libitum*. The generation and breeding described were approved by the Landesamt für Natur, Umwelt, und Verbraucherschutz Nordrhein-Westfalen (LANUV), Germany (animal applications: 84-02.04.2015.A405, 84-02.04.2018.A401 and 81-02.04.2021.A130).

### Embryo dissection

The embryos were dissected in cold 1x phosphate buffer saline (1x PBS; VWR, Radnor, PA, USA) with 0.4% bovine serum albumin (BSA; Sigma-Aldrich, St. Louis, MO, USA) (E6.5-E11.5) or in 1x PBS (E12.5-E15.5), and fixed in 4% paraformaldehyde (PFA; Carl Roth, Karlsruhe,Germany) overnight at 4°C. Embryos were then washed in 1x PBS, and either dehydrated in a methanol series to be used for *in situ* hybridization, or stored in 1x PBS at 4°C until further processing.

### Whole mount in situ hybridization

cDNA from E12.5 embryos was used as a template to amplify and generate the *Dkk4 in situ* hybridization probe of ∼0.7 Kb and covering the entire coding region (*Dkk4_ISH_forward: 5’- CTCCGAGAGACCAGAGTGAC-3’ and Dkk4_ISH_reverse: 5’-CACAACAACAAGTCCCGTGT-*

*3’)*. The PCR products were cloned into the pCRII dual promoter (T7 and SP6) vector (Invitrogen, Waltham, MA, USA) and the DIG-labeled probes were generated according to the manufacturer’s recommendations (Roche Applied Science, Indianapolis, IN, USA). *In situ* hybridization was performed on embryos from different stages per detailed published protocols (Wilkinson, 1998). The embryos were photographed using a DFC 450C camera fitted onto an M165 FC stereomicroscope (Leica Microsystems, Wetzlar, Germany).

### Immunofluorescence staining and imaging

For sections, embryos were washed with 1x PBS and cryoprotected in 30% sucrose (Sigma-Aldrich) in 1x PBS overnight at 4°C. After embedding in Tissue-Tek® (Optimal Cutting Temperature Compound, Sakura Finetek USA INC), the blocks were sectioned using a CM1850 Cryostat (Leica Biosystems) with an 8-10 µm thickness. Post-natal back skins were directly embedded in Tissue-Tek® and sectioned. For immunofluorescence staining, embryo sections were washed with washing buffer containing 0.2% Triton^TM^ X-100 (Sigma-Aldrich) in 1x PBS and blocked for 1 hour (h) in blocking buffer made of washing buffer and 1% heat-inactivated goat serum (HIGS) (Life Technologies, Waltham, MA, USA). Skin sections were fixed in 4% PFA for 10 min before they were washed and blocked in blocking buffer containing 10% HIGS. Embryo and skin sections were then incubated with the primary antibodies overnight at 4°C (LHX2, 1:1000, #3030529 Merck, Darmstadt, Germany; SOX9, 1:1000, #AB5535 EMD Millipore, Burlington, MA, USA; SOX2, 1:150, #14981182 Invitrogen), washed in washing buffer the next day, incubated with the secondary antibody and DAPI (1:1000; AppliChem, VWR) for 1 h at RT and mounted with Prolong™ Gold Antifade reagent (Cell Signaling Technology, Danvers, MA, USA). For whole-mount staining, embryos from E6.5 to E11.5 were fixed in 4% PFA for 2 h-overnight at 4°C, washed in 1x PBS, then in buffer containing 0.1- 0.3% Triton^TM^ X-100 and blocked with blocking buffer made of washing buffer with 10% serum. Primary antibody (Brachyury 1:200, #AF2085 R&D Systems, Minneapolis, MN, USA) and secondary antibody stainings were performed overnight at 4°C on two successive days. Embryos were washed 3 times for 30 min with washing buffer after each step and then mounted on 35 mm glass bottom dishes (Ibidi, Munich, Germany) using Vectashield® mounting medium (Vectorlabs, Newark, CA, USA) and embedded in 1% low melting agarose (Lonza, Basel, Switzerland). Images were obtained using a SP8 confocal microscope (Leica Microsystems) or a Meta 710 confocal microscope (Carl Zeiss AG, Oberkochen, Germany) and image acquisition was performed using LAS X (Leica Microsystems) or Zen lite (Carl Zeiss AG) software, respectively.

### Whole-embryo clearing

Embryos from E12.5 to E15.5 were cleared using ethyl cinnamate (Sigma-Aldrich) as previously described (Klingberg et al., 2017). Fixed embryos were washed three times for 10 min in 1x PBS and dehydrated in an ethanol series (50%, 70%, then twice in 100%). Every dehydration step was for 1 h at room temperature with gentle horizontal rotation. After dehydration, the embryos were transferred to ethyl cinnamate and imaged, when they became completely transparent, on an SP8 confocal microscope (Leica Microsystems).

### Image analyses

The number of LHX2- or SOX9- and EGFP-positive placodal cells was quantified through manual counting using ImageJ (Schneider et al., 2012). The graphs were generated using Prism (GraphPad, San Diego, CA, USA). Images of *Dkk4-Cre:mTmG/nTnG* embryos or sections were processed through ImageJ and Imaris (Bitplane, Belfast, United Kingdom).

### Reproducibility

Each experiment was repeated independently at least three times to ensure reproducibility.

### Data availability

The data, information, detailed protocols, reagents and mouse models in this study are available from the corresponding author upon reasonable request.

## Acknowledgement

We thank the CECAD *in vivo* research facility (Branko Zevnik) for generating and maintaining the mouse lines and the CECAD imaging facility for microscopy support. We are grateful to members of the lab and colleagues for critical reading of the manuscript. The work was partly funded by the Deutsche Forschungsgemeinschaft (DFG, German Research Foundation) - Project-ID 73111208 - SFB829 “Molecular Mechanisms regulating Skin Homeostasis”, Project A12 to H.B. The funders had no role in study design, data collection and analysis, decision to publish, or preparation of the manuscript.

## Author contributions

Conceptualization: H.K. and H.B.; Methodology: H.K. and H.B.; Software: H.K.; Formal Analysis: H.K.; Investigation: H.K. and H.B.; Writing: H.K. and H.B.; Visualization: H.K.; Supervision, Project administration and Funding Acquisition: H.B.

## Supplementary Figure Legends

**Supplementary Figure 1:**
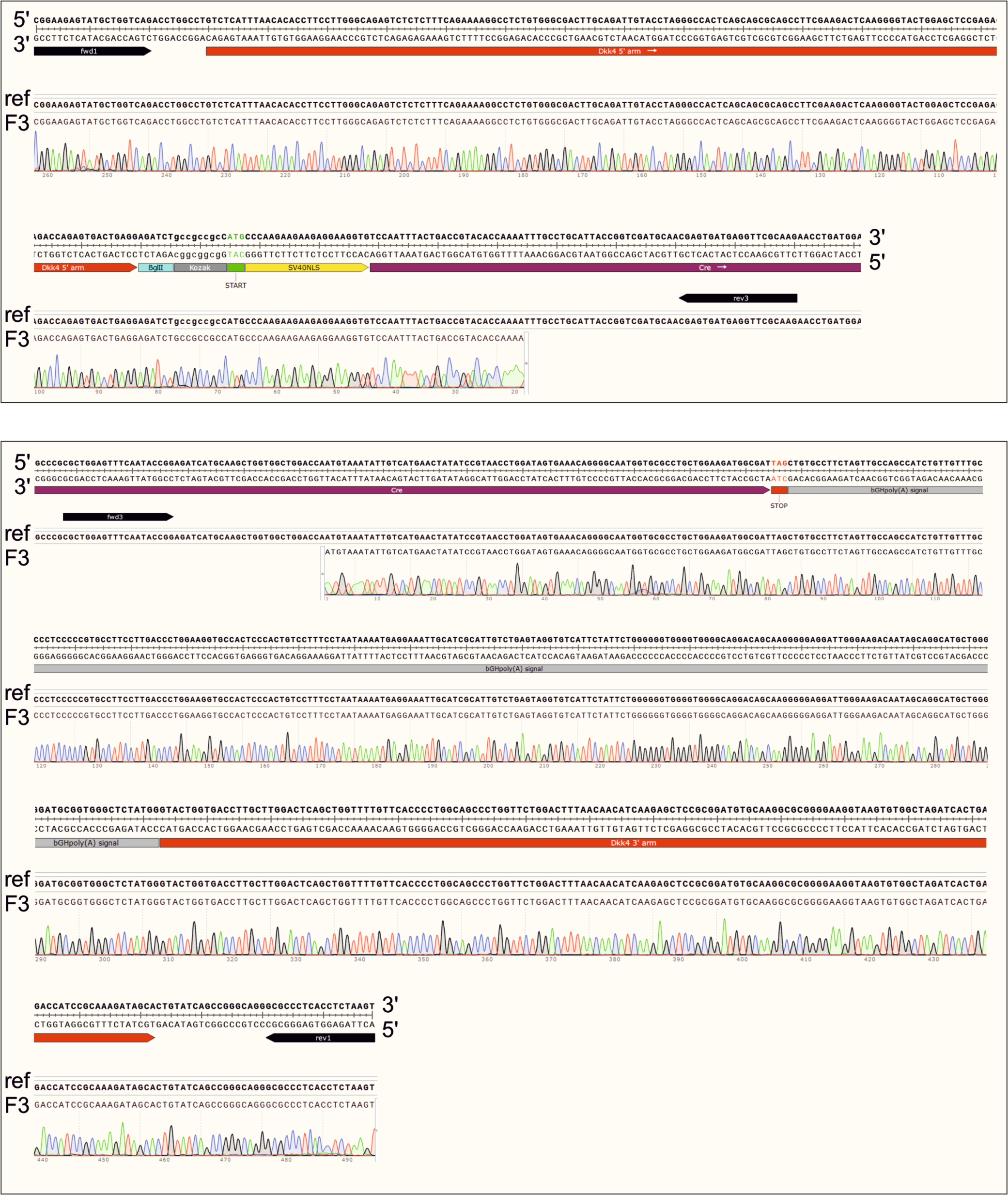
Sequencing and alignment of the 5’ and 3’ PCR amplifications of the targeted-locus in *Dkk4-Cre* Founder 3. PCR using (fwd1+ rev3, top) or (fwd3+rev1, bottom) was sequenced using the primer rev3 for the 5’-region and fwd3 for the 3’-region. The sequencing results of Founder 3 (F3) were aligned to the reference sequence (ref, ssDNA template) representing the insertion of the *Cre* template into the *Dkk4* locus (SnapGene®).

**Supplementary Figure 2:**
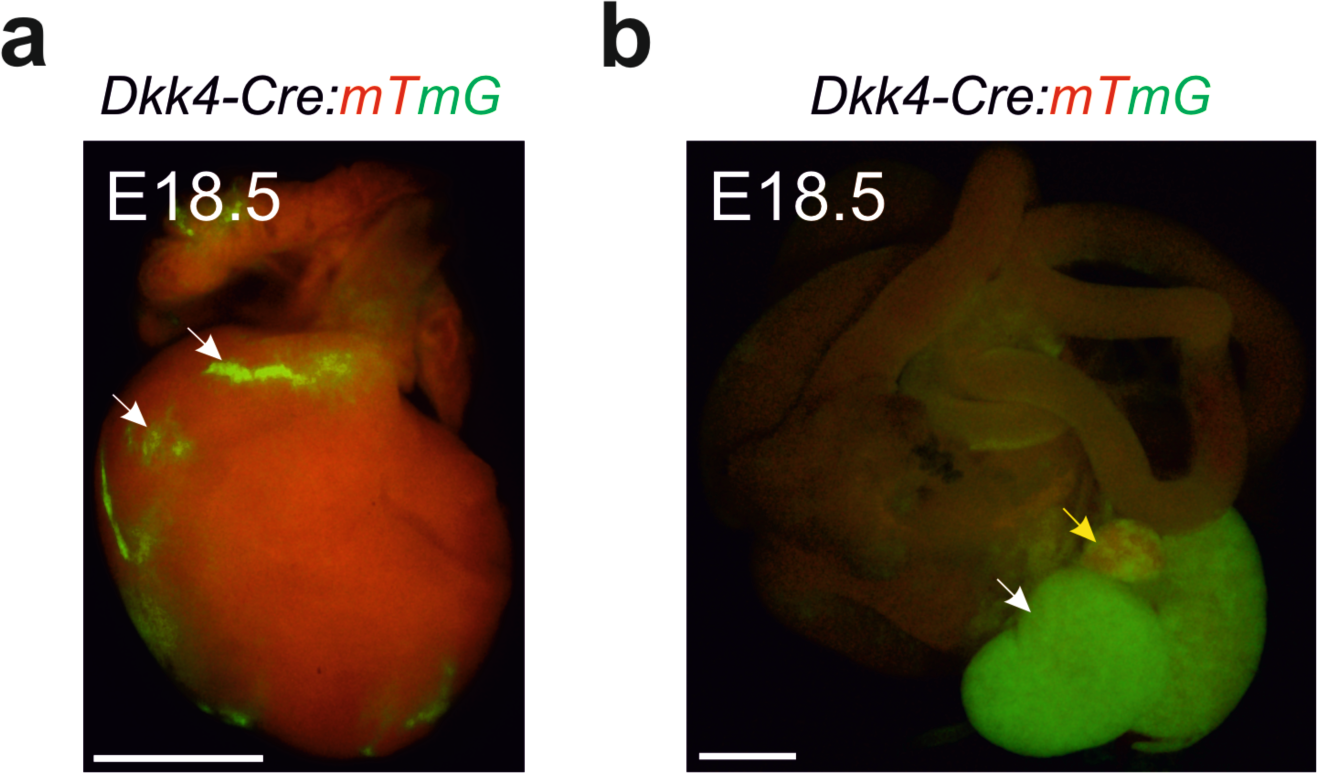
*Dkk4*-*Cre*-expressing lineage (EGFP) is detected in the heart, kidney and adrenal gland. Stereomicroscope images of EGFP-positive signals in the heart (white arrows in a), the kidney (white arrow in b) and adrenal gland (yellow arrow in b) from *Dkk4- Cre^KI/wt^:mTmG^tg/wt^* embryos at E18.5 (scale bar: 1000 µm).

**Supplementary Figure 3:**
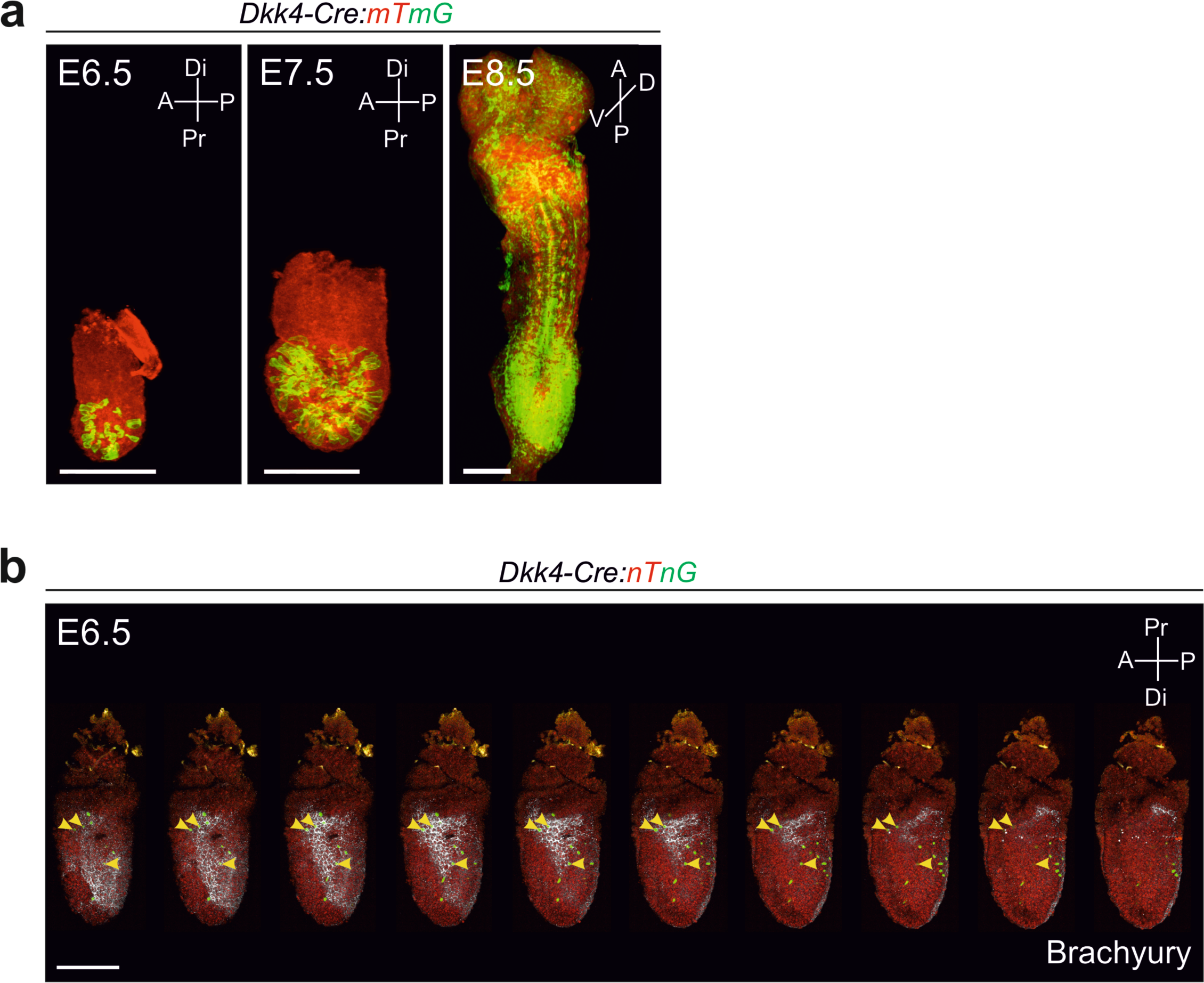
*Dkk4-Cre* expressing cells in early embryos. (a) Confocal images of E6.5 – E8.5 *Dkk4-Cre^KI/wt^:mTmG^tg/wt^*embryos with wide-spread EGFP-expression in the anterior and posterior body regions (scale bar: 200 µm). Embryos are orientated as indicated: Anterior (A), posterior (P), proximal (Pr), distal (Di), ventral (V) and dorsal (D). (b) Single confocal Z-plane images of an E6.5 *Dkk4-Cre^KI/wt^:nTnG^tg/wt^* embryo with individual EGFP-positive cells localized in the epiblast/primitive streak and mesoderm, some of which co-localized with Brachyury in white (yellow arrowheads) (scale bar: 200 µm). Axes of orientation as in (a).

